# A system-wide approach to monitor responses to synergistic BRAF and EGFR inhibition in colorectal cancer cells

**DOI:** 10.1101/194845

**Authors:** Anna Ressa, Evert Bosdriesz, Joep de Ligt, Sara Mainardi, Gianluca Maddalo, Anirudh Prahallad, Myrthe Jager, Lisanne de la Fonteijne, Martin Fitzpatrick, Stijn Groten, A. F. Maarten Altelaar, René Bernards, Edwin Cuppen, Lodewyk Wessels, Albert J. R. Heck

## Abstract

Intrinsic and/or acquired resistance represents one of the great challenges in targeted cancer therapy. A deeper understanding of the molecular biology of cancer has resulted in more efficient strategies, where one or multiple drugs are adopted in novel therapies to tackle resistance. This beneficial effect of using combination treatments has also been observed in colorectal cancer patients harboring the BRAF(V600E) mutation, whereby dual inhibition of BRAF(V600E) and EGFR increases antitumor activity. Notwithstanding this success, it is not clear whether this combination treatment is the only or most effective treatment to block intrinsic resistance to BRAF inhibitors. Here, we investigate molecular responses upon single and multi-target treatments, over time, using BRAF(V600E) mutant colorectal cancer cells as a model system. Through integration of transcriptomic, proteomic and phosphoproteomics data we obtain a comprehensive overview, revealing both known and novel responses. We primarily observe widespread upregulation of receptors tyrosine kinases and metabolic pathways upon BRAF inhibition. These findings point to mechanisms by which the drug-treated cells switch energy sources and enter a quiescent-like state as a defensive response, while additionally reactivating the MAPK pathway.

## Introduction

Despite the progressive development of novel drugs for personalized medicine, intrinsic and/or acquired resistance remains a major limitation of targeted anticancer therapies (Groenendijk & Bernards, 2014; Ahronian & Corcoran, 2017). Most of these drugs target components of the mitogen-activated protein kinase (MAPK) signaling pathway, which contains oncogenes such as KRAS, BRAF and the epidermal growth factor (EGFR) (Dhillon *et al*, 2007; Miyamoto *et al*, 2017). The use of monotherapy to inhibit these oncogenes has often been found to be ineffective due to reactivation of signaling pathways. For instance, upregulation of upstream components such as receptors tyrosine kinases (RTKs) in KRAS mutant lung and colorectal cancer (CRC) or of downstream components such as KRAS wild-type in CRC have been revealed to be responsible for intrinsic drug resistance (Sun *et al*, 2014; Karapetis *et al*, 2008).To overcome intrinsic and/or acquired resistance, combined drug treatments are frequently replacing single-agent targeted therapies (Komarova & Boland, 2013; Webster, 2016; Tong *et al*, 2017). An elegant example of bypassing intrinsic resistance using a multi-target approach has been demonstrated in BRAF(V600E) mutant CRC (Sundar *et al*, 2017). Whereas BRAF inhibitor (BRAF*i*) monotherapy is highly effective in BRAF(V600E) mutant melanoma, response rates in BRAF(V600E) mutant CRC are quite poor (Chapman *et al*, 2011; Kopetz *et al*, 2010). Multiple independent studies on CRC found a crucial role of EGFR as a key driver of resistance to BRAF*i* monotherapy (Prahallad *et al*, 2012; Corcoran *et al*, 2012; Klinger *et al*, 2013). In congruence with the role of EGFR in conferring resistance to BRAF*i*, the suppression of tyrosine phosphatase non-receptor type 11 (PTPN11) — which is required to transduce signals from EGFR and other RTKs to the downstream MAPK pathway — also sensitizes BRAF(V600E) CRC cells to BRAF inhibition (Prahallad *et al*, 2015). Consequently, the identification of EGFR as a mediator of intrinsic resistance to BRAF*i* in CRC has led to initiation of several clinical trials which combine inhibition of both EGFR and BRAF (BRAF*i*+EGFR*i*), or of other MAPK pathway members (Sundar *et al*, 2017).

Although the BRAF*i*+EGFR*i* combination treatment is more effective than BRAF*i* monotherapy in CRC (Kopetz *et al*, 2017), it remains unclear whether EGFR is the only or most potent synthetic lethal co-target of BRAF(V600E) in CRC. Addressing this issue requires an understanding of the cellular response to drug treatment across different molecular levels. Such multilayer approaches could elucidate different branches of the signaling network, and track how perturbations propagate to gene and protein expression in driving resistance. Several studies have already highlighted the widespread responses to drug treatment in cancer using multi-omics approaches and adequate data integration (Mertins *et al*, 2016; Zhang *et al*, 2016, 2014). Advances in next-generation sequencing and proteomics approaches (Altelaar *et al*,2012; Mathivanan *et al*, 2008) in combination with enhanced data integration solutions have paved the way for such important investigations (Nesvizhskii, 2014; Wang & Zhang, 2014). Notably, the integrated use of transcriptomics and (phospho)proteomics has recently demonstrated its power in describing physiopathological processes through phenotype and proteotype analysis (Faulkner *et al*, 2015; Low *et al*, 2013; Cutillas, 2015; Mertins *et al*, 2016).

In this study we analyze and integrate proteomics, phosphoproteomics, and transcriptomics data to follow the molecular responses over time upon perturbation with BRAF*i*, EGFR*i*, or their combination in CRC cell lines (**Figure 1**). We aim to study whether there are other post-translational or transcriptional mechanisms — in addition to EGFR and the MAPK pathway — that are activated upon treatment, in order to identify novel targets that may overcome innate and acquired resistance to BRAF inhibition.

**Figure 1.**
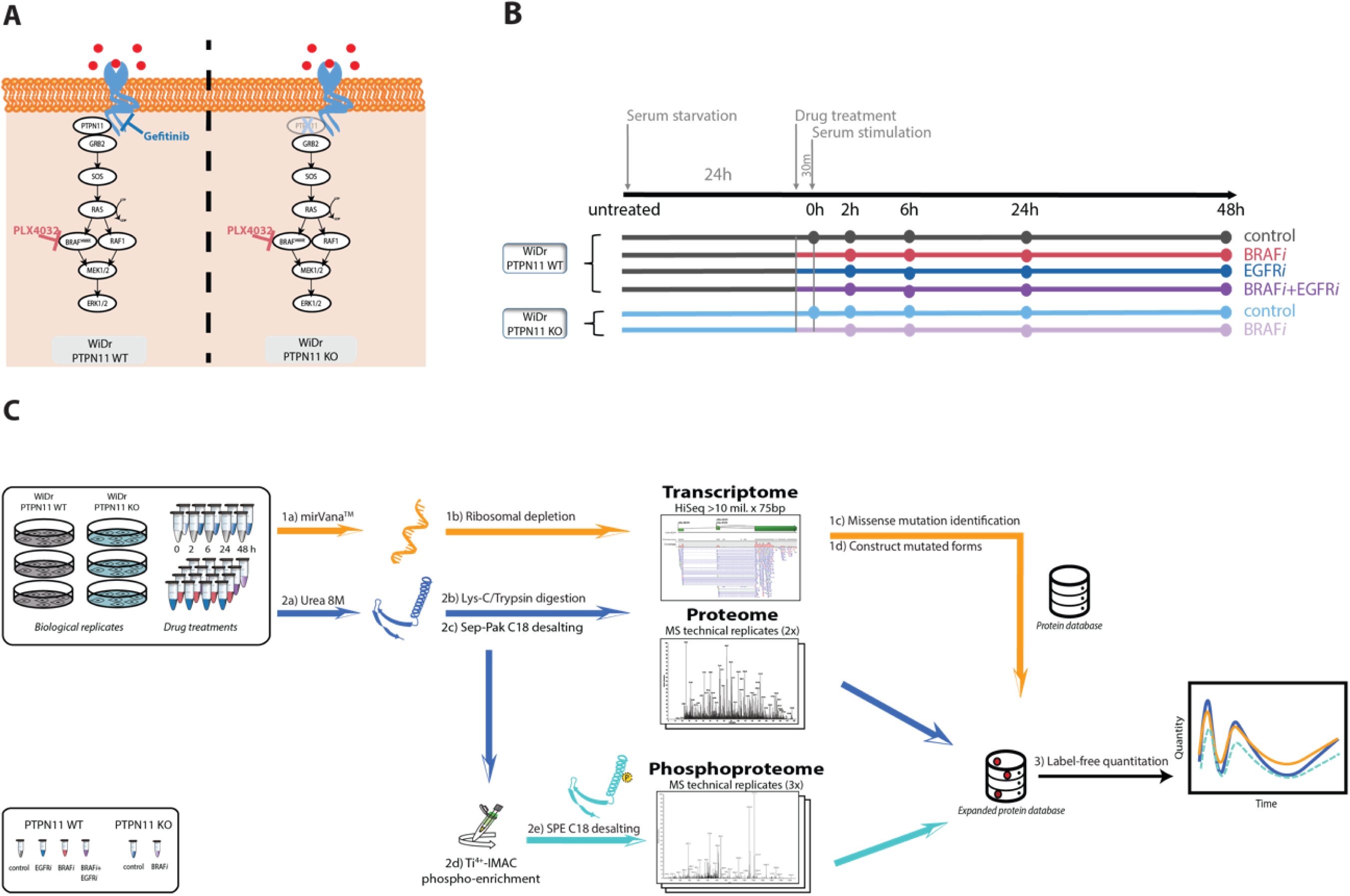
Overview of study design. **A. Biological model.** Schematic representation of the MAPK signaling pathway whereby BRAF(V600E), the drugs and KO target sites are highlighted. **B. Experimental design.** WiDr and WiDr PTPN11 KO cells were cultured for transcriptome, proteome and phosphoproteome analysis. For each of the six treatments the time-course of events is indicated **C. Workflow employed in this study.** For the transcriptomics analysis, mRNA libraries were prepared and sequenced using Illumina NextSeq. Quantitation in proteomics and phosphoproteomics analysis was done by label-free quantitation. Phosphoproteomics was performed after Ti^4+^-IMAC phosphopeptide enrichment. For the integration of transcriptome, proteome and phosphoproteome approach, a customized protein database was developed in oder to include identified missense mutations for peptide identification and quantification.

## Results

### A multi-omics overview of BRAF mutated colorectal cancer cell response to targeted drug treatment

The WiDr CRC cell line harboring the BRAF(V600E) mutation was selected as a model system for our analyses (Noguchi *et al*, 1979; Chen *et al*, 1987). To study the differences in signaling we treated the WiDr cells with either BRAF*i* or EGFR*i* monotherapies or with the combination treatment BRAF*i*+EGFR*i*. Additionally, we employed a WiDr PTPN11 knockout (KO) cell line treated with BRAF*i* (BRAF*i* in PTPN11 KO) to investigate if there are functional differences between PTPN11 KO and EGFR*i*. Cell growth was synchronized by serum starvation for 24 hours (h), followed by 30 minutes (min) incubation with or without drugs before serum stimulation (Supplementary Methods). Unstimulated control and PTPN11 KO control cells were immediately harvested — indicated as T=0 h throughout this study — whereas stimulated samples were collected in a time course at 2, 6, 24, and 48 h after treatment (**Figure 1B**). We performed transcriptomic (RNA-seq) and (phospho)proteomic profiling at each of the time points as indicated in **Figure 1C**. This study was designed to capture the initial responses to the different drug treatments and the elicited RTK signaling, monitoring their effects on gene and protein expression, and the onset of feedback mechanisms. For this approach, a customized protein sequence database was generated using RNA-seq data to account for WiDr-specific non-synonymous variants (**Figure 1C** and Supplementary Methods).

The phosphorylation profile of ERK (MAPK1) (**Figure 2A,** top panel) was used as a positive control to verify the drug-induced regulation and overall quality of the label-free quantitative (phospho)proteomics approach. In concordance with previous studies (Prahallad *et al*, 2015), pERK becomes downregulated upon BRAF*i*, and this effect is enhanced by the addition of EGFR*i*, reaching, as expected, the same levels as of the BRAF*i* in PTPN11 KO cell line. Complementary Western blots (**Figure 2A,** bottom panel) show excellent correlation with the label-free phosphoproteomics quantified data. Further quality analysis demonstrates high correlation between respective biological replicates for each omics dataset, with median correlations of 0.99, 0.93 and 0.83 for the transcriptomics, proteomics and phosphoproteomics data respectively (**Figure 2B**). Quantified proteomics data points show, as expected, slightly higher variability in comparison to RNA-seq data (Haider & Pal, 2013), while phosphoproteomics data points exhibit even higher variability. The final (phospho)proteomics dataset consisted of 5692 quantified protein groups and 7141 quantified Class I phosphosites (localization probability > 0.75), both of which were measured in at least one timepoint of any of the six applied conditions (**Figure S1**). The transcriptome dataset contained a final list of 21,446 genes. We developed a Graphical User Interface (GUI) to facilitate rapid data comparison. The GUI enables selection of a specific gene to immediately visualize a comparison of its expression profile at transcriptomic and (phospho)proteomic levels (**Figure 2C**).

**Figure 2.**
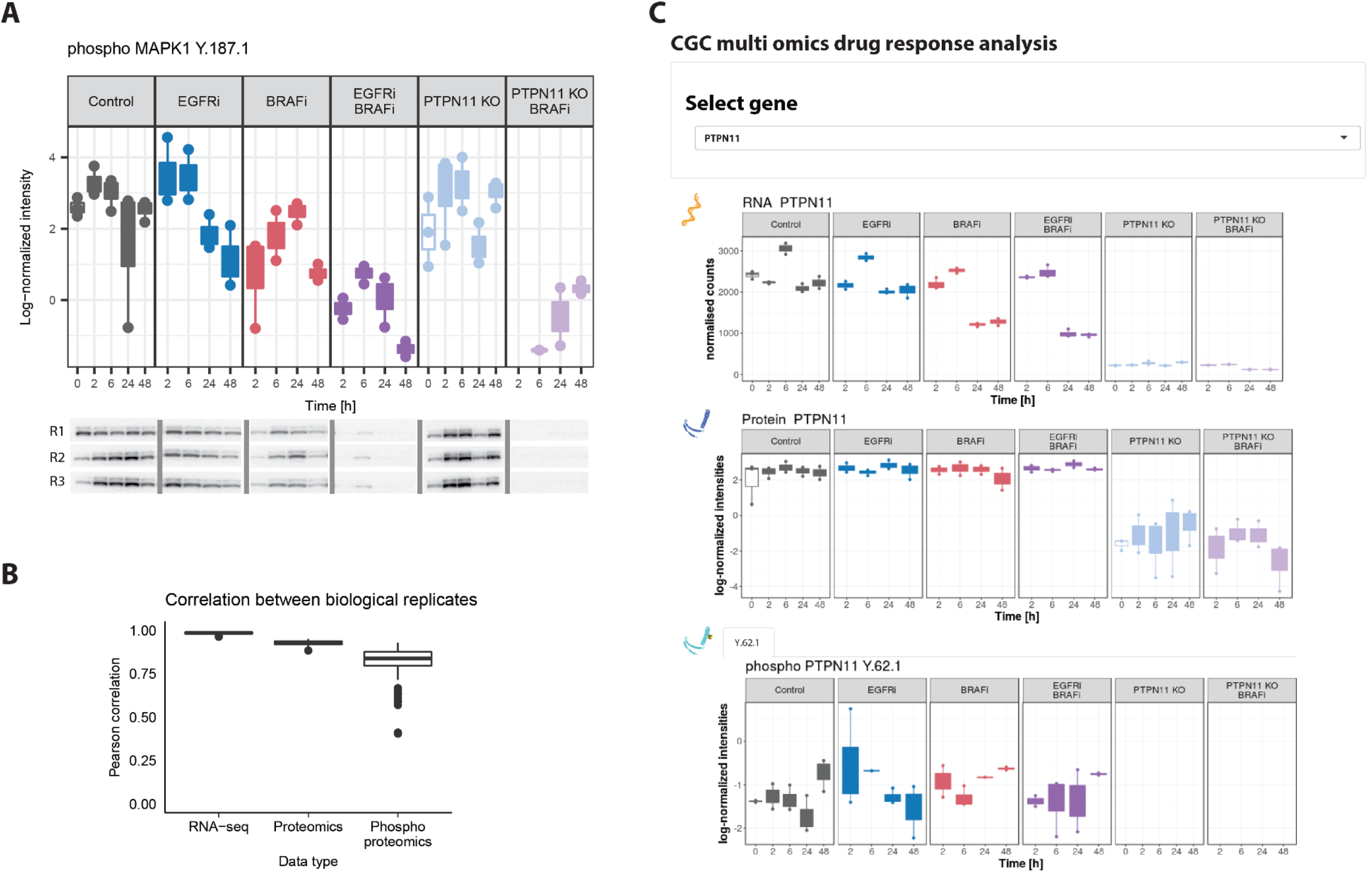
High data quality enables reliable overview of gene-level response across omics datasets. **A. Label-free phosphoproteomics and phosphoWB provide alike regulation patterns.** The LC-MS/MS quality control was assessed by quantification of ERK phosphorylation (top panel) in all three biological replicates (R1, R2 and R3), and subsequent validation was done via Western blots (bottom panel). **B. Quality analysis of reproducibility at each omics level.** Replicate consistency was assessed by inter-replicate Pearson correlation. In the phosphoproteomic analysis, selection of technical replicates involved discarding the poorest replicates before quantification. The resulting processed data were further filtered to only include data that was quantified in at least one timepoint of any condition. Analysis indicates good correlation (R>0.8) among the three biological replicates for all three omics datasets. **C. Cross-omics Graphical User Interface output.** The GUI enables exploration of the multi-omics data for specific genes under all the tested conditions. Users can select a measured gene of interest from a dropdown menu, and visualize complete data at the protein, phosphoprotein and transcript level, over the time-course and over all experimental conditions. An example output for the gene CDK1 is shown. The user can toggle which number of sites is seen, but not the exact phoshposite.

To extract a more general overview of the effect of the different experiments on the cells, we performed Principal Component Analyses on each omics data type (**Figure 3A**). The trend along the first two principal components is similar for all data types. Principal Component 1 (PC1) represents the variation in the measurements over time in the BRAF*i* treated samples (BRAF*i*, BRAF*i*+EGFR*i*, and BRAF*i* in PTPN11 KO). This variation is slightly greater in BRAF*i*+EGFR*i* and BRAF*i* in PTPN11 KO samples compared to the samples treated with BRAF*i* only. Notably, the onset of the variation in the direction of PC1 occurs earlier in the transcriptomic data (after 6 h) than in the proteomics data (after 24 h) reflecting the delay from translation to transcription. Principal Component 2 (PC2) reflects the variation in the measurements of the analysed cells in non-BRAF*i* treated samples (control, EGFR*i* and PTPN11 KO control). In the proteomics data, PC2 also clearly captures the variation induced by the PTPN11 KO.

**Figure 3.**
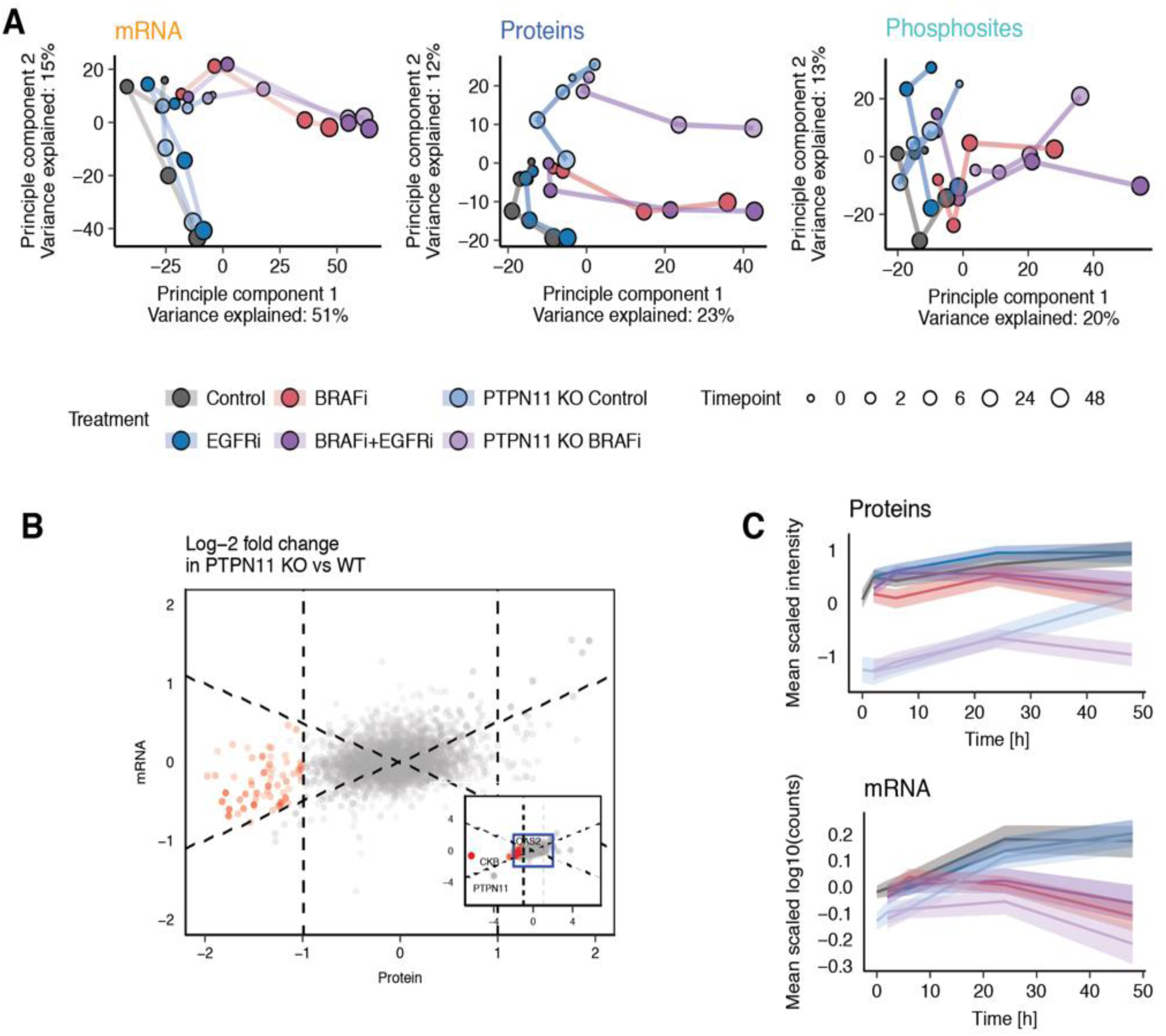
PCA analysis highlights similarities and differences between drug treatments and omics data types. **A.** Principal Component Analysis (PCA) of transcriptomics, proteomics and phosphoproteomics data elucidates similar global trends in the data. **B.** Scatterplot of log_2_-fold changes of protein (x-axis) and mRNA (y-axis) levels in PTPN11 KO compared to PTPN11 WT cells. **C.** Average expression profiles of the 65 proteins (red dots in B) that are downregulated at the protein but not at the mRNA level. The expression of each mRNA/protein is mean-centered. The solid lines indicate mean expression of the 65 mRNA/proteins and the shaded area indicates the 95 % confidence interval of the mean.

### PTPN11 knockout induces post-transcriptional downregulation

To further investigate the segregation of the isogenic cell line PTPN11 KO from PTPN11 wild-type (WT) as observed predominantly at the protein level, we performed differential protein and mRNA expression analyses using a linear model (limma), where the PTPN11 status (KO and WT), the drug treatments (BRAF*i*, EGFR*i* and BRAF*i*+EGFR*i*) and the time-points (0, 2, 6, 24 and 48 h) were set as coefficients. Our aim was to elucidate how the PTPN11 status affects proteins and mRNAs differently by contrasting differential expression of proteins and mRNAs (**Table S1**).

At the protein level, we observe the strongest difference in creatine kinases brain-type (CKB),with a log_2_-fold-change of -6.3 in PTPN11 KO cell line compared to PTPN11 WT. Remarkably, CKB shows minimal difference on average in mRNA expression (log_2_-fold-change = -0.65) (**Figure S2A**). Interestingly, CKB has been recently considered responsible for promoting survival in WiDr CRC cells, by regulating the reservoir of high-energy phosphates in tissues to sustain intracellular energetic requirements (Loo *et al*, 2015). Further analysis identified 64 additional proteins that have significant differential protein expression (FDR = 0.1) between PTPN11 KO and PTPN11 WT, a large negative log_2_-fold-change of < -1 at the protein level, and a small log_2_-fold-change at the mRNA level (< protein log_2_ fold-change/2) (**Figure 3B**).

These 65 proteins also demonstrate a consistent time-course profile in which expression at both the protein level and the mRNA level is initially upregulated in all six experimental conditions (**Figure 3C**). At T > 24 h, downregulation of both the mRNA and protein level occurs in BRAF*i* treated conditions, whereas expression levels in other conditions are upregulated or remain unchanged. This observation suggests that these 65 proteins can be regulated at the mRNA expression level, despite downregulation in an mRNA-expression independent manner upon PTPN11 KO. The consistency in expression profiles across the 65 proteins also suggests that these genes are to some extent co-regulated.

We next investigated whether these 65 proteins are functionally related by performing an enrichment analysis using the MSigDB Hallmarks gene-sets (Liberzon *et al*, 2015). Our analysis reveals a strong enrichment of the interferon alpha (IFN-α) and gamma (IFN-γ) response gene-sets (enrichment > 15-fold, *p* < 10^-9^, **Table S1**) which are known to suppress cell viability through the JAK/STAT pathway (Ivashkiv & Donlin, 2013). Interestingly, PTPN11 negatively regulates the INF-induced JAK/STAT pathway by dephosphorylating STAT1 on both residue Y701 and S727 (Du *et al*, 2005). In line with these observation, STAT1 is also significantly downregulated at the protein level in the PTPN11 KO cells (*p* < 10^-12^) in our dataset and shows a time-course profile similar to the aforementioned 65 genes (**Figure S2B**).

### System-wide propagation of drug perturbation

A key aspect of our study was to provide a system-wide understanding of the propagation of cellular responses from signaling (phosphoproteomics) to gene transcription (transcriptomics) and then translation and protein expression (proteomics) in response to BRAF and/or EGFR inhibition. We therefore performed correlation-based hierarchical clustering on the 1500 phosphosites, mRNAs and proteins exhibiting the highest variance within each dataset, which resulted in eight clusters for each omics dataset (**Figure S3**). We observe that in the transcript-, phosphosite- and protein clusters, all BRAF*i* treated samples (BRAF*i*, BRAF*i*+EGFR*i* and BRAF*i* in PTPN11 KO) exhibit similar clustering profiles distinct from non-BRAF*i* treated samples.

Next, we investigated the biological function of all the clusters by performing an enrichment analysis based on transcription factor-target gene (transcriptomic clusters) and kinase-substrate (phosphoproteomics cluster) relationships. We also conducted enrichment analyses using the hallmarks gene-sets from MSigBDg (Liberzon *et al*, 2015) (**Table S2**): Two clusters, corresponding to an early treatment response within 2-6 h and late treatment response after 24-48 h in both the gene expression (transcriptomic) level and the signalling (phosphoproteomic) level are dowregulated upon BRAF*i* treated samples, and had a clear biological interpretation based on the enrichment analyses (**Figure 4A** and **4B**). We also distinguished four additional clusters characterized by phosphosites and transcripts, which show distinct upregulation after 2 h upon BRAF*i* treatment.

**Figure 4.**
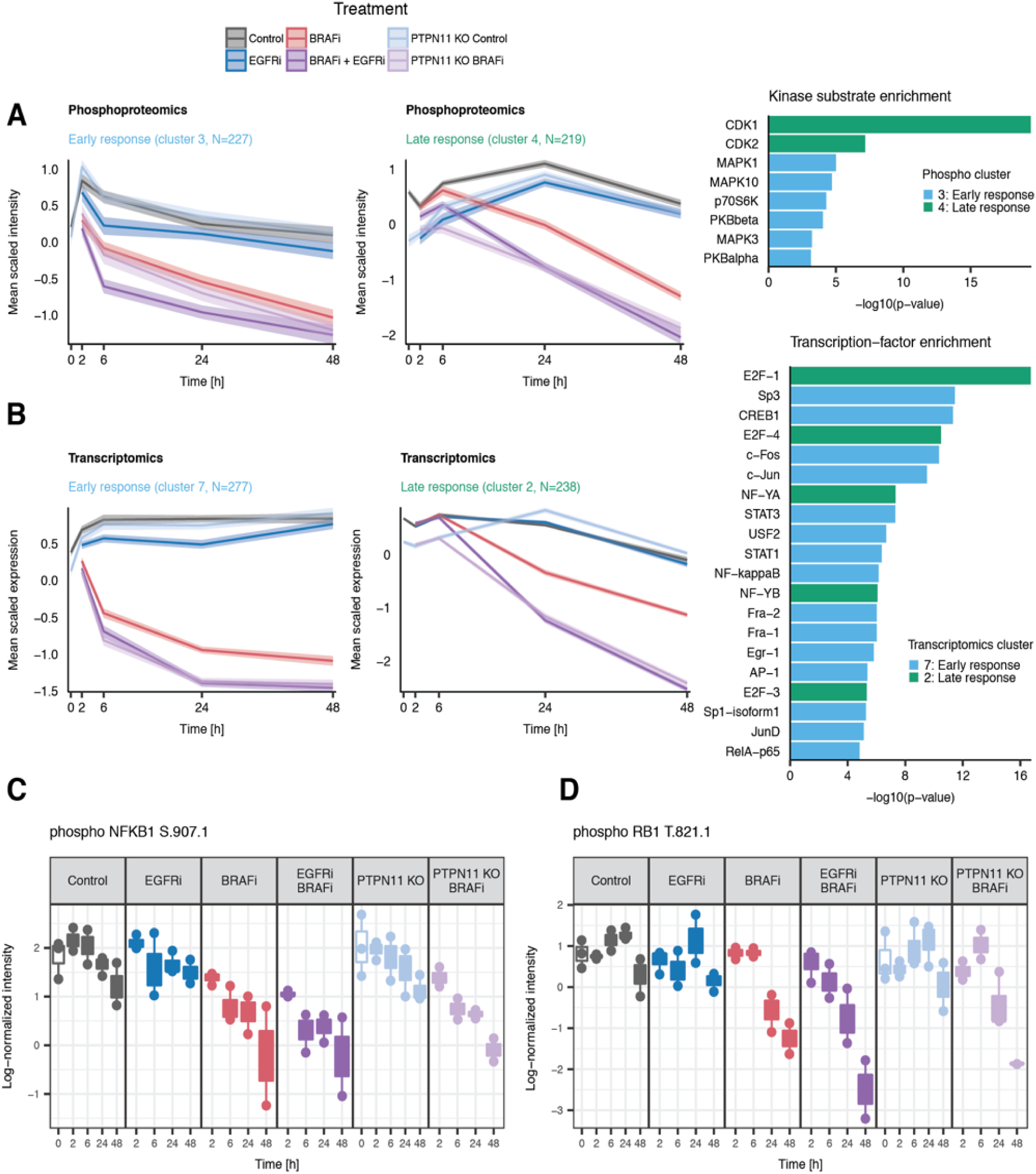
Clustering reveals a MAPK mediated early and CDK mediated late response to BRAF inhibition. **A.** Selected phosphoproteomics clusters show an early and late response of phosphosites in which phosphorylation decreases upon BRAF inhibition within 2-6 h (left panel) and after 24-48 h (middle panel). Enrichment analysis (right panel) indicates that the early response cluster is enriched for substrates of kinases located downstream of BRAF in theMAPK pathway, whereas the late response cluster is enriched for substrates of cyclin-dependent kinases. **B.** Selected transcriptomics clusters show a similar early (left panel) and late (middle panel) response. Enrichment analysis (right panel) reveals that the early response cluster is enriched for target genes of transcription factors that are downstream of the MAPK pathway, and the late response cluster is enriched for targets of the E2F-transcription factor. **C.** Deactivation of NF-κB is a consequence of MAPK pathway deactivation as shown by downregulation of phosphosite S907 on NFKB1 exclusively in all the BRAFi treated samples. **D.** Downregulation of RB1 at phosphosite T821 inhibits E2F1 activity and induces cell cycle arrest at late timepoints in BRAFi treated cells.

#### MAPK, PI3K-AKT and NF-κB signaling are involved in the early response

The early response phosphoproteomics cluster is enriched for substrates of kinases belonging to the MAPK and PI3K-AKT pathways (**Figure 4A**, right panel). An immediate decrease in phosphorylation of MAPK1 (ERK2) and MAPK3 (ERK1) substrates occurs upon BRAF inhibition, which is consistent with dephosphorylation of Y187 and T185 phosphosites on MAPK1 and Y204 on MAPK3 (**Figure 2A** and **S4A**). The early response cluster also includes p70S6K (RPS6KB1) substrates. Similarly, this is consistent with the dephosphorylation of the kinase itself (at the S427 phosphosite) after 2-6 h upon BRAF*i* and BRAF*i*+EGFR*i* (**Figure S4B)**. PKBα (AKT1) and PKBβ (AKT2) substrates are also enriched in this cluster exhibiting an unexpected deactivation of AKT upon BRAF*i* monotherapy with respect to what was previously shown (Prahallad *et al*, 2012). The extent of deactivation could not directly be resolved since we did not detect the relevant phosphosites of AKT1 and AKT2 in our dataset. Therefore, we confirmed deactivation of AKT by Western blot (**Figure S7**). Consistent with the early deactivation of MAPK and AKT signaling observed in the phosphoproteomics cluster, the transcriptomics early response cluster is enriched for targets of transcription factors downstream of the MAPK and AKT pathways (**Figure 4B**, right panel), such as CREB1, FOS, JUN, STAT1, STAT3 and MYC. The early response cluster is absent in the proteomics data (**Figure S3B**), possibly reflecting a time delay between transcription and translation.

Surprisingly, NF-κB, which is not typically considered to be downstream of the MAPK pathway (Moynagh, 2005), is amongst the transcription factors most enhanced in the early response cluster. The phosphorylation profile of S907 on NFKB1 is part of the early response cluster (**Figure 4C**). NFKB1 S907 phosphorylation induces activity (Cartwright *et al*, 2016), suggesting that gene expression of the early response cluster is partially driven by inactivation of NF-κB. Our proteomics data show no substantial variation in the protein expression of NFKB1 under any experimental condition (**Figure S4C**), indicating that activity of NFKB1 is likely inhibited by dephosphorylation. The residue S907 of NFKB1 is a possible substrate of GSK3β kinase (Demarchi *et al*, 2003), which is downstream of MAPK in the pathway, and GSK3β substrates are enriched in the early response phosphoproteomics cluster (albeit not significantly after multiple testing correction). This observation suggests that deactivation of NF-κB is a consequence of the observed MAPK pathway deactivation.

#### Drug treatment affects the cell cycle and cell proliferation at the late cellular response

The late response cluster reflects the effect of drug treatment on cell cycle and proliferation, both at the transcriptomics and phosphoproteomics level. The phosphoproteomics late response cluster is highly enriched for substrates of the cell cycle regulators CDK1 and CDK2 (**Figure 4A**, right panel). This finding is corroborated by decreased CDK1 (**Figure S4D**) and CDK2 (**Figure S4E**) protein expression. Similarly, the transcriptomic late response cluster is enriched for target genes of the cell cycle regulators E2F1, E2F4, and E2F3 (**Figure 4B**, right panel). The connection between late response phospho-signaling and gene expression is mediated by downregulation of the phospho-residue T821 on RB1 (**Figure 4D**), a substrate of CDK2, while total RB1 protein expression remains relatively constant (**Figure S4F**). This downregulation induces binding of RB1 to E2F1 thereby inhibiting E2F1 activity (Lentine *et al*, 2012), and promoting cell cycle arrest (Henley & Dick, 2012) under BRAF*i* conditions at the late timepoints. Further enrichment analysis of the MSigDB hallmarks gene-sets revealed enrichment for proliferation-related gene sets including E2F targets, genes involed in the G2M checkpoint, and mitotic spindle genes (**Table S2**).

We also observe a late response cluster in the proteomics data, which is strongly enriched for targets of MYC and E2F (proteomics cluster 4). MYC and E2F targets are also enriched in the transcriptome early response cluster, again indicative of the delay between transcription and translation.

#### Combined BRAFi and EGFRi treatment leads to a distinct metabolic response

In addition to the early and late response clusters, we observe a set of transcripts, proteins and phosphosite clusters that are upregulated in BRAF*i* treated samples compared to control and EGFR*i*-only treated samples. These clusters include phosphoproteomics cluster 5, transcriptomics clusters 5 and 6, and proteomics cluster 2 (**Figure S3**) and exhibit the strongest regulation at 48 h. mRNA and protein clusters in this set were enriched for metabolic processes including peroxisome, fatty acid and bile acid metabolism (**Table S2**). Notably, mRNA cluster 6 is enriched for targets of the transcription factors C/EBPβ, Sp1, HNF-1α and HNF-4α, which are involved in fatty acid metabolism and in the gluconeogenesis pathway (Desvergne *et al*, 2006). To pinpoint in more detail which biological processes are upregulated in the BRAF*i+*EGFR*i* treated samples at 48 h, we performed differential expression and enrichment analysis of transcripts and proteins that were significantly upregulated at this time point with respect to control (log_2_-fold-change > 1, FDR < 0.05). KEGG and Reactome pathway analyses revealed significant upregulation in metabolic processes and pathways including the pentose phosphate pathway (PPP), the TCA cycle and the lipid metabolism (**Table S3**). Taken together, these observations suggest that the cells reprogram to a distinctive metabolic pattern in response to the treatment.

Similarly, cluster 5 of phosphoproteomics data was enriched in substrates of the calcium/calmodulin dependent protein kinase II isoforms (CAMKIIα/β/γ/ δ) and of pyruvate dehydrogenase kinase isozyme 1 (PDHK1). Besides being responsible for Ca^2+^ homeostasis under pathophysiological conditions (Anderson, 2011), CAMKII is also linked to resistance to apoptosis due to its metabolic activation caused by increased levels of acetil-CoA and intermediates in the PPP (McCoy *et al*, 2013; Huang *et al*, 2014). PDHK1 rather regulates glucose and mitochondrial metabolism by phosphorylating pyruvate dehydrogenase E1 component subunit alpha (PDHA1) on serine residues (Harris *et al*, 2002; Dupuy *et al*, 2015). In our data, PDHA1 expression is constant at the protein level in the BRAF*i*+EGFR*i* samples at 48 h (**Figure S5C**), while phosphorylation is upregulated at S232 on PDHA1 (**Figure S5A**) but downregulated at S293 (**Figure S5B**). As PDHK1 is not detected at the protein level but only at mRNA level (**Figure S5D**, **S5E** and **S5F**), we are limited in our interpretation with respect to PDHA1 modulation. Conversely, our proteomics data demonstrate an activated TCA cycle upon BRAF*i*+EGFR*i* as evidenced by significant upregulation of the TCA cycle enzymes IDH1, IDH2, SUCLG1 and SUCLG2 (log_2_-fold-change > 1, FDR < 0.05). The observed enrichment of peroxisomal proteins, indicated by upregulation of CPT2, SLC25A20, SLC25A1, SLC25A10 and SLC25A13, suggests fatty acids may be used as energy source under the drug treatment inducing stress conditions (Röhrig & Schulze, 2016) (**Figure S6**). Taken together, our omics data suggest that upon combined BRAF*i*+EGFR*i* treatment, WiDr CRC cells increase their mitochondrial oxidative activity through fatty acid synthesis and uptake, possibly as a defensive response.

### Feedback responses aim to reactivate the MAPK pathway

Cells are able to preserve a stable homeostatic balance through activation of feedback mechanisms in a counteractive manner as a response to perturbation. Here, we sought to elucidate the molecular response to drug treatment by investigating the presence of potential feedback response mechanisms. In general, a homeostatic response to pathway stimulation can be either downregulation of a signal transducer or upregulation of a signal inhibitor. To study this systematically, Legewie *et al.* collected gene expression profiles of responses to pathway stimulation (Legewie *et al*, 2008). By plotting the expression changes of known signal transducers and inhibitors of major signaling pathways (MAPK, PI3K, cAMP, TGFβ, JAK/STAT) against their mRNA half-lives, they observed a striking pattern. All responding genes where a) signal inhibitors and b) short-lived. They called these short-lived signal inhibitors Rapid Feedback Inhibitors (RFIs), which, according to them, constitute a fast and efficient homeostatic response mechanism. We here investigated if a similar design principle applies to pathway inhibition, plotting the log_2_-fold change mRNA expression after 2 h — compared to control T=0 h in each condition — against the mRNA half-lives obtained by Legewie *et al.* (**Figure 5A**). We observe four interesting features in this plot. Firstly, all responding genes (DUSP1, DUSP4, DUSP6, DUSP8, DUSP10 and SPRY1) are short-lived signal inhibitors, consistent with the observations of Legewie *et al.* Secondly, only genes regulating MAPK pathway signaling are responding, suggesting that no other signaling pathways are affected by either growth-factor stimulation or BRAF*i*. Thirdly, all samples show strong upregulation of DUSP1, DUSP8 and DUSP10; presumably in response to the serum stimulation at T=0 h. Finally, only BRAF*i* treated samples exhibit strong downregulation of DUSP4, DUSP6, and SPRY1, demonstrating that RFIs can also be downregulated in response to pathway inhibition, in an attempt to counteract pathway inhibition.

**Figure 5.**
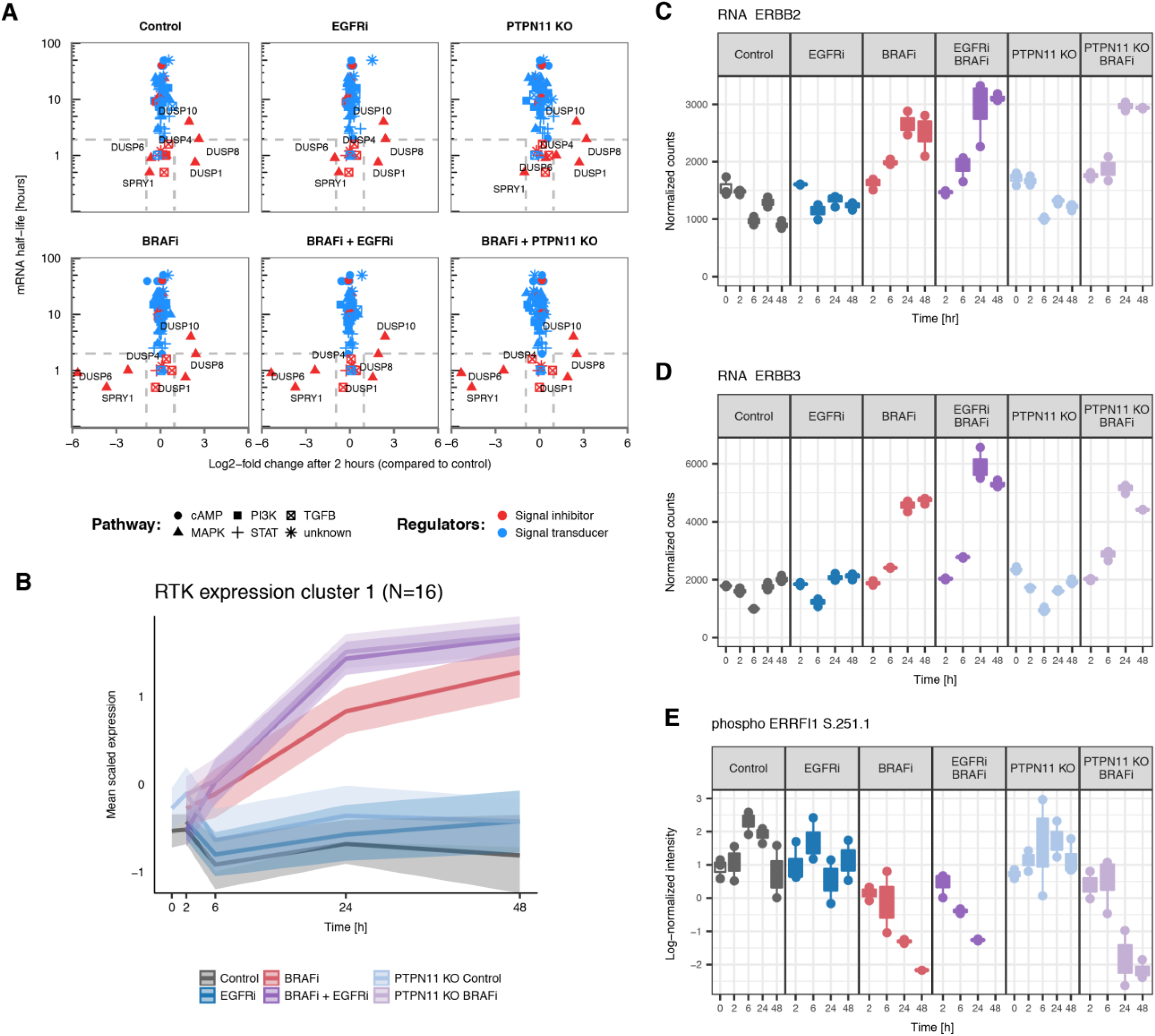
Feedback mechanisms aimed at restoring MAPK signaling activity. **A.** Log_2_-fold-change of mRNA expression plotted against mRNA half-life of regulators of major signaling pathways. Only short-lived negative regulators of MAPK signaling respond to growth-factor stimulation or BRAF inhibition. In all conditions, DUSP1, DUSP8 and DUSP10 are upregulated in response to serum stimulation at T=0 h. DUSP4, DUSP6 and SPRY1 are downregulated only in BRAFi treated samples, in response to BRAF inhibition. **B.** Scaled mRNA expression levels of a cluster of RTKs. 18 out of the 35 RTKs are upregulated. **C.** and **D.** mRNA expression of ERBB2 and ERBB3 **E.** Phosphorylation expression of the negative regulator ERRFI1 is downregulated at residue S251 in BRAFi samples.

Next, we examined the various RTKs that were upregulated at the transcriptome level upon BRAF inhibition. We performed hierarchical clustering of the mRNA expression of all RTKs based on Pearson correlation to minimize bias. Our transcriptome data reveal strong RTK upregulation upon BRAF inhibition from the mid-timepoint (T=6-24 h) onward. Of the 35 RTKs expressed in WiDr cells, 16 were upregulated (**Figure 5B**), including ERBB2 and ERBB3 (**Figure 5C and D**). Protein expressions levels of selected targets were further verified by Western blots performed on the same WiDr and WiDr PTPN11 KO lysates used for omic analysis (**Figure S7**). ERBB2 and ERBB3 are known to confer acquired resistance mechanisms as evidenced by reduced response to EGFR and BRAF(V600E) inhibitors (Montero-Conde *et al*,2013; Sun *et al*, 2014; Jain *et al*, 2010; Temraz *et al*, 2016). Interestingly, in addition to the ERBB2/ERBB3 upregulation, the regulator ERBB receptor feedback inhibitor 1 (ERRFI1), which interferes with ERBB family member homo- and hetero dimer formation (Liu *et al*, 2012), is downregulated in all BRAF*i* treated samples (**Figure 5E**). Altogether, our data suggests the existence of an additional mechanism through which WiDr CRC cells may activate ERBB signaling to compensate for MAPK pathway inhibition.

### Combined BRAFi and EGFRi treatment induces a stable but reversible growth arrest

To further investigate the response of BRAF(V600E) mutant cells to combined BRAF*i*+EGFR*i* treatment, we exposed WiDr cells to the combination of both drugs for a prolonged period of 78 days, after which few surviving cells were still present in small colonies that did not further expand, indicative of growth arrest (**Figure 6A**, panel 1). To elucidate whether the arrested cells retained the ability to resume their cell cycle upon drug withdrawal, we switched the cells to conventional medium (“drug off”). As shown in **Figure 6A** (panel 3), 6 days after drug withdrawal the cells start to grow again and reached confluency within 10 days (**Figure 6A**, panel 4). Similar results are obtained when WiDr cells were subjected to BRAF*i*+EGFR*i* treatment during 5 days (**Figure 6B**, panel 1) and subsequently switched to conventional medium (**Figure 6B**, panel 2, 3 and 4). Altogether, our data suggest that combined BRAF and EGFR inhibition in WiDr BRAF(V600E) mutant cells induces an incomplete apoptotic response, resulting in surviving cells temporarily arrested but with retained proliferative capacity. This observation could be relevant for translating combined BRAF*i*+EGFR*i* treatment into the clinical practice. In this regard, elucidating the cell cycle stage in which these cells are arrested may contribute to a better understanding of how to induce complete cell death.

**Figure 6.**
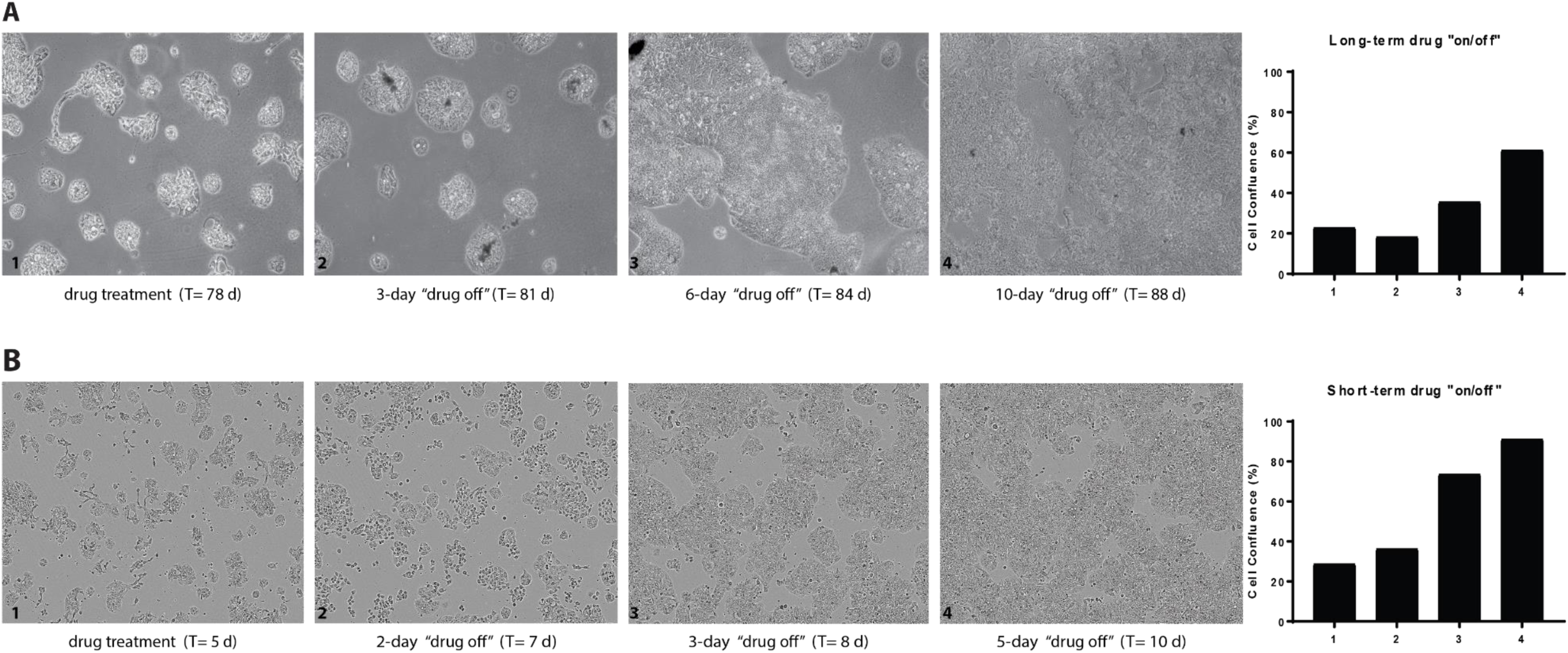
Combination treatment induces temporary cell cycle arrest. **A.** Prolonged exposure to BRAF*i*+EGFR*i* treatment causes the appearance of resistant colonies in WiDr cell line. Interruption of drug treatment (“drug off”) induces cells to resume proliferation. **B.** A shorter period of “drug off” (72 h) further confirms the presence of reversibly arrested WiDr cells by showing a rapid resumption of cell proliferation after “drug on” (5 d). The difference in confluence between long- and short-term may depend on the ability of cells to leave thequiescent adaptive status after continuous treatment.

### ERBB inhibitors provide limited benefit in BRAF(V600E) CRC treatment

Next we sought to determine if the observed upregulation of ERBB2 and ERBB3 upon BRAF*i*+EGFR*i* treatment could be further exploited. We first studied whether inhibition of ERBB2 and ERBB3 in combination with BRAF*i* and EGFR*i* may lead to complete cell death by using gefitinib, lapatinib and sapitinib as known tyrosine-kinase inhibitors of EGFR, EGFR/ERBB2 and EGFR/ERBB2/ERBB3, respectively. Next, we explored if there was an optimal drug concentration, which would be synergistic with the maximum tolerated dose of PLX4032 (BRAF*i*) (Supplementary Methods). As expected (Prahallad *et al*, 2012), WiDr CRC cells are resistant to monotherapy of either gefitinib or lapatinib or sapitinib, with decreasing viability only at very high, i.e. cytotoxic, concentrations (**Figure 7A**). All three viability curves depict a 60 % decrease in cell viability upon addition of 3 μM PLX4032, with limited benefit from the combination with EGFR*i*. We did not observe a significant difference in growth inhibition across the different double treatments, suggesting that additional inhibition of ERBB2 and ERBB3 does not provide further synergy with BRAF*i*.

**Figure 7.**
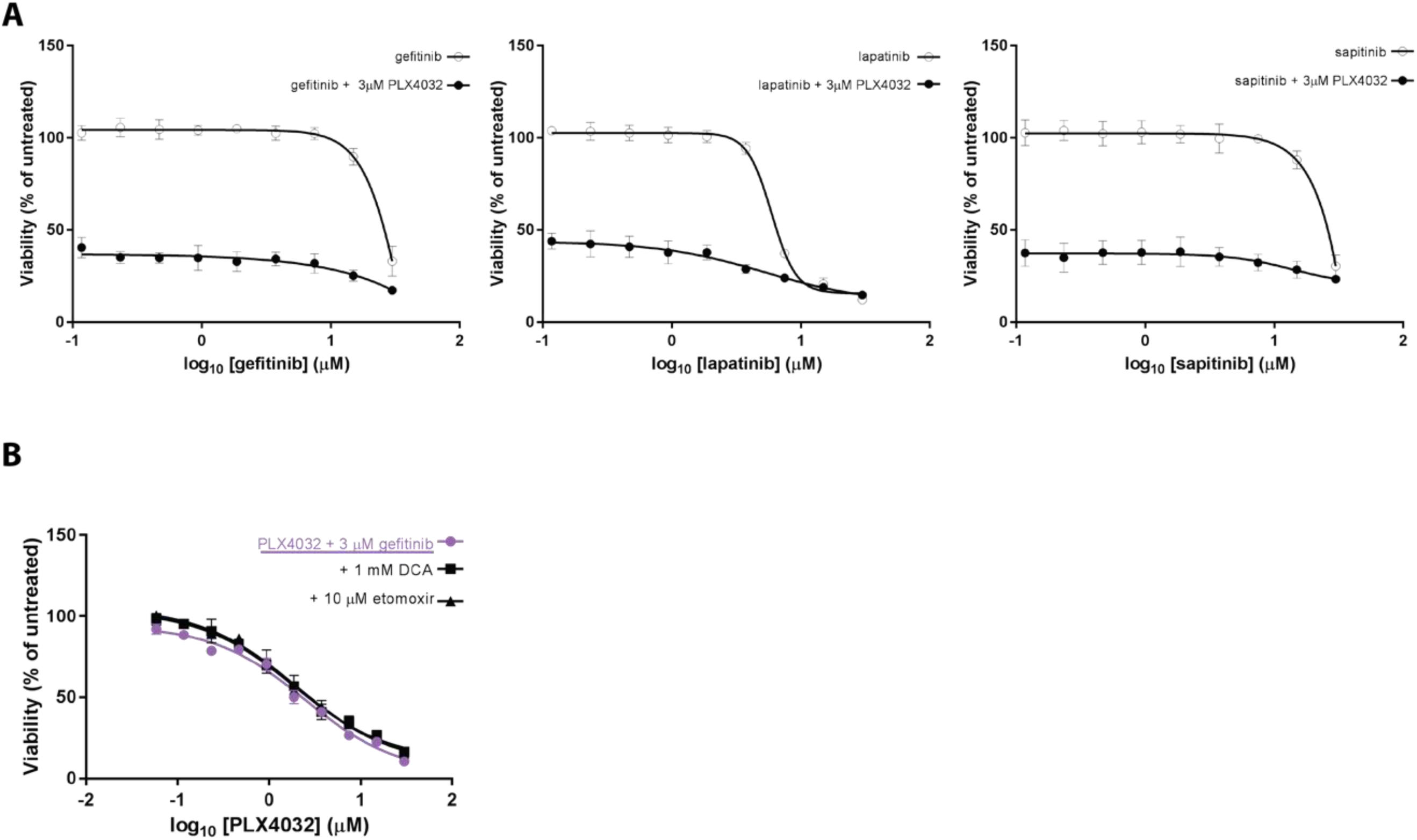
Assessment of WiDr CRC cell growth by combination treatments of BRAFi and ERBB or metabolic inhibitors. **A.** Comparison of mono- and double therapy on WiDr CRC cells growth. All three graphs show inhibition of either EGFR, EGFR/ERBB2 or EGFR/ERBB2/ERBB3 is un-effective as a monotherapy. Moreover, concomitant inhibition of ERBB2 and ERBB3 does not provide further benefit to the synergistic effect of BRAF(V600E) and EGFR inhibitors. **B.** WiDr cell confluence is measured comparing double and triple treatments. The addition of etomoxir or DCA as third metabolic inhibitors does not show additional benefit to the BRAF*i*+EGFR*i* treatment.

### Inhibitors of metabolic enzymes provide limited benefit in BRAF(V600E) CRC treatment

We further evaluated combination treatments to target MAPK pathway together with the TCA cycle or with the fatty acid β-oxidation. For this porpuse we used two readily available metabolic drugs: etomoxir, a CPT1 inhibitor, and dichloroacetate (DCA), a pan-inhibitor of PDKs, to determine if inhibition of CPT1 or PDKs enhances sensitivity to therapy with PLX4032 and gefitinib (EGFR*i*). CPT1 is directly upstream of CPT2 in the peroxisomal fatty acid oxidation processes and is responsible for transporting long chain fatty acids from the cytosol to the mitochondrial matrix (Lodhi & Semenkovich, 2014; Pucci *et al*, 2016). CPT2 is significantly upregulated in our experiments, nevertheless, to the best of our knowledge, a direct inhibitor of CPT2 has not been previously reported. PDKs are responsible for deactivation of PDHA1 through phosphorylation of serine residues in PDHA1 (Harris *et al*, 2002; Dupuy *et al*, 2015) (**Figure S5**). We hypothesized that inhibition of PDKs can increase the mitochondrial oxidative state and consequently the amount of reactive oxidative species (ROS) in the cytoplasm causing apoptosis due to high toxicity. We therefore evaluated the combination treatment of each metabolic inhibitor (etomoxir and DCA) with BRAF*i* and EGFR*i* on WiDr cell viability. After assessing IC50 of both metabolic drugs (**Figure S8A**), we selected the clinical doses of 10 μM etomoxir (Ito *et al*, 2012; Holubarsch *et al*, 2007; Samudio *et al*, 2010) or 1 mM DCA (Dunbar *et al*, 2014; Fox *et al*, 1996) and we performed two dose-response curves in the presence of 3 μM gefitinib and increasing concentrations of PLX4032. The first curve was obtained by adding the metabolic drug simultaneously to the BRAF*i* and EGFR*i* (T=0 h) (**Figure S8B**), and the second by adding it 96 h after BRAF*i* and EGFR*i* (T=96 h) (**Figure 7B**), when metabolism is supposed to be significantly upregulated according to our data. In both cases, we do not observe any significant differences in the viability curves of the triple treatments in comparison to the double treatments. These findings suggest that inhibition of CPT1 or PDKs does not increase sensitivity to therapy with PLX4032 and gefitinib.

## Discussion

In this study we performed an integrative quantitative multi-omics analysis to obtain a system-wide molecular characterization of signaling perturbation over time in WiDr CRC and WiDr PTPN11 KO cell lines, after drug inhibition targeting either BRAF(V600E) and/or of EGFR. Our data reveal that all samples treated with BRAF*i* show similar response in both PTPN11 WT and KO, with a less pronounced effect following BRAF*i*-monotherapy. This indicates that main signalling responses depend on the inhibition of BRAF(V600E), whereby the additional inhibition of EGFR by drug further amplifies the effect. Additionally, EGFR*i*-only treated cells exhibit similar responses to PTPN11 KO samples, confirming that suppression of this secondary signaling pathway confers sensitivity to BRAF*i* in CRC (Prahallad *et al*, 2015).

By comparing proteomics and transcriptomics data, we identified a set of genes that are exclusively downregulated at the protein level, upon PTPN11 KO. These proteins are negative regulators of the interferon pathway (Porritt & Hertzog, 2015), involved in controlling immune response. Downregulation of negative regulators may support the immune response elicited by PTPN11 *in vivo* (Mainardi et al., unpublished data). This finding might be relevant for the development of therapeutic approaches that aim to inhibit PTPN11 activity.

The integrated omics analysis enabled us to trace the system-wide drug response upon treatment, from signaling – e.g. inactivation of kinases downstream of the MAPK pathway - through transcription – e.g. inhibition of genes downstream in the MAPK pathway. Shutting down MAPK signaling results in downregulation of CDK signaling, inducing cell cycle arrest at a later stage. Besides the inactivation of the MAPK pathway, all three datasets show an increase of oxidative metabolic processes, with significant upregulation of enzymes involved in lipid metabolism and the TCA cycle. We further observed that treatment with BRAF*i* induces upregulation of RTKs, including ERBB2 and ERBB3, which was found to be more pronounced when co-treated with EGFR*i* or in PTPN11 KO cells.

A subset of cells survived the BRAF*i*+EGFR*i* treatment and could start to proliferate again after drug removal. This indicates that these cells survive by utilizing different metabolic regimes, pointing at potential future avenues on how to target these cells. However, in our work combining inhibition of the MAPK pathway and specific metabolic processes did not result in any significant difference in cell viability. The precise protective role of metabolic adaptation in the ability of cells to tolerate drug treatment remains elusive, and further studies are required.

Apart from the upregulation of metabolic processes, all adaptive responses we observed appear to be homeostatic responses that try to re-activate the MAPK pathway, but their attempting seem to fail in the range time of this study. Despite apparent activation of ERBB family members, we do not find an additional benefit of inhibiting ERBB2 and/or ERBB3 in combination with EGFR*i* and BRAF*i*, suggesting that these homeostatic responses are not necessarily functional under the tested conditions. Further studies are required to establish the more general implications of these findings, investigating different cell lines and several medium conditions that more closely mimic physiological environments. Importantly, we do not find any evidence of parallel signaling pathways being activated in response to drug treatment.

Together, this suggests that reactivation of the MAPK pathway is a requirement for BRAF(V600E) mutation CRC cells to become resistant to BRAF inhibition. This view is supported by observations in the clinic that resistance to MAPK pathway inhibitors is typically mediated by mutations or amplification in the MAPK pathway in CRC patients (Ahronian *et al*, 2015).

Our findings on metabolic rewiring do not show any direct impact when targeted, on the time scale measured in this study, but might still be relevant to therapy as two recent studies in melanoma demonstrated the dependence of resistant cells on mitochondrial respiration (Corazao-Rozas *et al*, 2013; Cesi *et al*, 2017). These studies highlight the importance of studying the complete omics landscape to identify relevant drug targets. The integrative multi omics approach employed here provides a time based and in depth view of the signaling mechanisms involved in drug response. Our findings highlight the importance of measuring these different levels simultaneously as exemplified by the RTKs regulation and PTPN11 specific signals. We expect to enable and accelerate future research into these mechanisms by making the data resource available.

## Materials and Methods

### Experimental design

Colorectal tumor cell line WiDr and WiDr PTPN11 KO were used as model to study resistance upon drug treatment. Cells were grown in three biological replicates. Cells were starved for 24 h and then treated with 3 μM PLX4032 (BRAF*i,* BRAF*i* in PTPN11 KO), 3 μM gefitinib (EGFR*i*) or combination of the two inhibitors (BRAF*i*+EGFR*i*). Unstimulated cells were used as control (T=0 h). Following EGFR stimulation cells were harvested at time points: 2, 6, 24 and 48 h after treatment. mRNA and protein lysates were extracted with *mir*Vana total RNA isolation kit and in urea 8 M buffer, respectively. The reported driver mutations (BRAF V600E, PIK3CA P449T, TP53 R273H) (Ahmed *et al*, 2013) were verified and the ploidy of the chromosomes checked against previous characterizations of WiDr/HT-29 (Chen *et al*, 1987) (**Figure S9**). Single nucleotide variants were called by GATK v3.4-46 (McKenna *et al*, 2010), copy number was determined using CONTROL-FREEC v10.4 (Boeva *et al*, 2012).

### (Phospho)proteomics

Cell lysates were first digested, then desalted and finally analyzed via mass spectrometry. Phosphopeptides were enriched using Ti^4+^-IMAC (Zhou *et al*, 2013) and subsequently desalted. A total of 312 samples were processed and 411 LC-MS/MS runs were collected for both phosphoproteome and proteome analysis. LC-MS/MS runs consisted of 2 h and 3 h gradient for proteome and phosphoproteome analyses, respectively. At least two technical replicates were collected for the proteome and three for the phosphoproteome analyses. Each dataset was checked for a preliminary quality control analysis, and phospho-technical replicates with poor correlation value (R^2^<0.7) were discarded. Finally, 383 raw files were selected for Label Free Quantification (LFQ) in MaxQuant (version 1.5.2.8). A customized database integrating variants with a predicted missense effect was used for (phospho)proteomics quantification. Output were processed with a Python package (PaDuA), generating the final dataset with 5692 protein groups and 7141 phosphosites Class I (phosphorylation localization probability ≥0.75). For complete information see Supplementary Methods.

### Transcriptomics

RNA integrity was checked on a Bioanalyzer 2100. All samples had a RNA integrity number (RIN) value ≥ 9.5. After RNA purification, libraries were prepared using a ribosomal depletion protocol (Illumina Truseq Stranded Total RNA kit with Ribo-Zero Human/Mouse/Rat). Libraries were sequenced on Illumina NextSeq to an average of 9.9 (±3.5) million reads per sample. From these RNA depleted RNA-seq libraries, 7.9 (±4.9) million reads mapped against the human reference genome (hg19), of which 36.6% (±8.6) correspond to mRNA regions. For complete information see Supplementary Methods.

### Western Blots

For both quality control and validation analysis, lysed cells were resolved by SDS-PAGE using NuPAGE Gel Electrophoresis Systems (Thermo Scientific). Protein detection was performed using Clarity ECL Western Blotting substrate (Bio-Rad) and blot imaging was performed using the Chemidoc Touch Imaging System (Bio-Rad). For complete information see Supplementary Methods.

### Differential expression analysis

All differential expression analyses were performed using limma (Ritchie *et al*, 2015). mRNA read counts were first transformed using voom (Gentleman *et al*, 2004). For the comparison of BRAF*i*+EGFR*i* to control samples, a linear model was fitted for each gene/protein/phosphosite with each condition (treatment and time-point pair) as a separate coefficient. The contrast between the BRAF*i*+EGFR*i* and control samples at 48 h conditions was used for enrichment analysis. For the comparison of the PTPN11 KO cell line to the PTPN11 WT cell line, a linear model was fitted for each gene/protein using cell line, time-point and treatment as coefficients,and subsequently contrasting the cell line coefficient. Because PTPN11 KO and EGFR*i* treatment are expected to have a similar biological effect, the treatment coefficient of PTPN11 KO controls and EGFR*i*-only treated samples were equated. Similarly, the treatment coefficient of PTPN11 KO cell line treated with BRAF*i* and BRAF*i*+EGFR*i* samples were equated.

### Clustering

Hierarchical clustering was performed on the 1500 phosphosites, proteins, or mRNAs with the highest variance over all conditions. The pairwise distance was calculated based on the Pearson correlation. The obtained distances were used for hierarchical clustering and the resulting tree was split into 8 groups for each dataset. Enrichment analysis were done using Fisher’s exact test. KinomeXplorer (Horn *et al*, 2014) was used for predicted kinase-substrate relations in phosphoproteomics clusters. TransFac (Matys *et al*, 2006; Wingender *et al*, 1996) database was used for transcription factor-target gene relations in transcriptomics clusters. Enrichment analysis of biological processes was done using the hallmarks gene sets from MSigDB (Liberzon *et al*, 2015). For complete information see Supplementary Methods.

### Cell proliferation assays

WiDr cells were seeded in 96-well plates (5000 cells/well) and 24 h later treated with: either gefitinib, lapatinib or sapitinib as single drug therapy (120 nM-30 μM) or in combination with 3μM PLX4032; double treatment of PLX4032 (30 nM-30 μM) with 3 μM gefitinib; triple treatment of PLX4032 (30 nM-30 μM) with 3 μM gefitinib and 10 μM etomoxir; triple treatment of PLX4032 (30 nM-30 μM) with 3 μM gefitinib and 1 mM DCA. Cellular growth was monitored in the IncuCyte ZOOM live cell microscope (Essen BioScience) and images were taken in phase contrast every 2 h until 7 days. For the IC50 of single treatments, WiDr cells were seeded in 96-well plates (10,000 cells/well) and 24 h later stimulated with either PLX4032 or etomoxir or DCA in the concentration range 30 nM-30 μM, 200 μM-0.2 μM, 100 mM-0.1 mM, respectively. Cellular growth was monitored in the IncuCyte ZOOM live cell microscope and images were taken in phase contrast every 2 h until 72 h. All proliferation assays were performed in four biological replicates. For complete information see Supplementary Methods.

### Drug off assays

For long-term assay, WiDr cells were seeded in one 6-well plate (200,000 cells/well), starved in serum-free media for 24 h, and then treated with 3 μM PLX4032 and 3 μM gefitinib in complete medium. Treatment was interrupted after 78 days for a “drug off” period of 10 days. Pictures were acquired using an ECLIPSE Ti-e inverted microscope (Nikon, Tokyo, Japan) at a magnification of 10x. For short-term assay, WiDr cells were seeded in one 96-well plate (5000 cells/well) and 24 h after treated with 3 μM PLX4032 and 3 μM gefitinib. Treatment was interrupted after 5 days for a “drug off” period of 5 days. Cell growth was monitored in the IncuCyte™ automated microscope (Essen Bioscience, Ann Arbor, USA) and phase-contrast images were collected every 2 h using objective Nikon 10x. For complete information see Supplementary Methods.

## Data and code availability

The mass spectrometry proteomics data have been deposited to the ProteomeXchange Consortium via the PRIDE (Vizcaíno *et al*, 2016) partner repository with the dataset identifier PXD007740. Sequence data has been deposited at the European Genome-phenome Archive (EGA) (Lappalainen *et al*, 2015), which is hosted by the EBI and the CRG, under accession number EGAS00001002654. Source code for all statistical analyses is available at https://bitbucket.org/evertbosdriesz/cgc-multi-omics. The Graphical User Interface (GUI) to visualize gene/protein/phosphosite expressions is available at https://cancergenomics.shinyapps.io/CGC_MultiOmics/. FAIR data portal is available at https://cgc.fair-dtls.surf-hosted.nl/demonstrator/.

## Acknowledgements

This work was conducted under the framework of the Gravity Program CGC.nl, funded by The Netherlands Organisation for Scientific Research (NWO). AR, MA, AJRH were further supported by the Roadmap Initiative Proteins@Work (project number 184.032.201), funded by The Netherlands Organisation for Scientific Research (NWO). We would also like to thank the PRIDE Team for assistance.

## Author contributions

AR, EB, JL, SM, GM and AP contributed to the design of the experiments. AR, JL, SM, GM, AP, MJ and LF and SG performed the experiments. AR, EB, JL and MF designed the data analysis. AR, EB, JL and MF performed the analysis. AR and EB wrote the manuscript.

## Conflict of interest

The authors declare that they have no conflict of interest.

## Supplementary Figures

**Figure S1. Data overview and quality control. A.** Phosphoenrichment performed on samples of each of the six culture conditions shows values of reproducible identifications (i.e. ∼80 % across all treatments). **B.** Summary table indicating the number of quantified proteins, phosphosites, and transcripts.

**Figure S2. Examples of differential regulation at the mRNA and protein level upon WiDr PTPN11 KO.** Examples of genes identified by the differential analysis between PTPN11 WT and PTPN11 KO, being downregulated at the protein level (top panel), but not mRNA level (bottom panel): **A.** Creatine Kinase B (CKB), **B.** Signal transducer and activator of transcription 1 (STAT1).

**Figure S3. Correlation clustering at all three omics levels.** A distance measure based on the Pearson correlation was used to generate 8 clusters for the phosphoproteomics (**A**), proteomics (**B**) and transcriptomics (**C**) data. MAPK and NF-κB regulated processes are enriched in clusters 3 (A) and 7 (C). Interferon alpha and gamma response genes, and targets of the transcription factors IRF-1, STAT1 are enriched in cluster 6 (B) and cluster 1 (C). Metabolic processes are enriched in clusters 5 (A), 2 (B) and 5 and 6 (C). PTPN11 KO is enriched in cluster 8 (C) specifically for RE1 Silencing Transcription Factor (REST1) targets.

**Figure S4. Expression profiles found to be enriched in specific clusters**. Early response clusters exhibit immediate downregulation of MAPK and PI3K-AKT pathways as shown by decreased phosphorylation of key residues after 2-6 h upon treatment: **A.** MAPK3 Y204 and **B.** RPS6KB1 S247. **C.** The unchanged protein expression of NFKB1 over all tested conditions and over time confirms the specific downregulation of its phosphosite S907. Late response clusters exhibit downregulation of the CDK pathway as further indicated by its downregulation at the protein level at the late stage: **D.** CDK1 and **E.** CDK2. **F.** The RB1 protein remains at constant expression levels at the protein level.

**Figure S5. Proteins involved in energy metabolism.** PDHA1 is a key metabolic enzyme, whose activity is negatively regulated by PDKs. Upon BRAF*i*+EGFR*i* treatment at 48 h, PDHA1 shows opposite pattern of phosphorylation levels on its residues **A.** S232 and **B.** S293; **C.** PDHA1 total protein expression, instead, remains constant. mRNA level of three out of four PDKs are shown: **D.** PDK1, **E.** PDK2 and **F.** PDK4. Opposite to PDK1 profile, both PDK2 and PDK4 show significant upregulation of their transcripts at 24 and 48 h.

**Figure S6. Observed changes in metabolic processes in the BRAF(V600E) cells upon combined BRAF*i* and EGFR*i* treatment.** Enrichment analysis revealed the TCA cycle, the peroxisome and lipid metabolism among the most upregulated processes upon BRAF*i*+EGFR*i* treatment at 48 h. Upregulation of the TCA cycle may depend on activity of PDHA1 whichcatalizes oxidative decarboxylation of pyruvate and on fatty acid β-oxidation. These processes could be used from WiDr cells as a defensive mechanism to generate energy during their quiescent state, and to prevent them from a complete apoptosis. The use of additional metabolic drugs targeting these processes may enhance the effect of BRAF*i* and EGFR*i* treatment. For example, indirect inhibition of PDHA1 (via PDKs) by DCA (red block) might increase the TCA cycle activity leading to toxic concentration of reactive oxygen species (ROS). Direct inhibition of CPT1, instead, might reduce TCA cycle activity and consequently reducing the energy needed for cell survival.

**Figure S7. Western blots validation on multiple protein and phospho-specific targets.** Validation of selected target regulations by Western blots analysis on the same WiDr PTPN11 WT (**A**, R2; **B**, R3) and WiDr PTPN11 KO (**C,** R2; **D**, R3) lysates used in our accross omics analysis. EGFR, ERK, P90 proteins and phosphoproteins are involved in the MAPK pathway, whereas AKT and p-AKT are involved in PI3K-AKT signaling. ERBB2 and ERBB3 were identified in RTK clustering together with INSR. IGF1R expression was also measured as it is thought to be closely related to INSR. The data showed consistently the reduction of p-EGFR, p-ERBB2, p-ERBB3 and p-IGF1R at 48 h in the combined BRAF*i* and EGFR*i* treatments. Nevertheless, ERBB2 and ERBB3 showed a higher expression at the latest time point in the WiDr cells concurrent with a decrease in EGFR.

**Figure S8. Cell proliferation assays for drug treatment. A.** Cell proliferation curves of WiDr cells upon treatments with either PLX4032 (BRAF*i*) or etomoxir (CPT1 inhibitor) or DAC (PDKs inhibitor) show similar profiles. Both three curves confirm WiDr cell line being resistant to each monotherapy. **B.** Cell proliferation curves of WiDr cells upon triple combination treatment. A fixed concentration of a metabolic drug (DCA or etomoxir) is added to PLX4032 (BRAF*i*) and 3μM gefitinib (EGFR*i*) at the same time (T=0 h). Resulting viability curves show same pattern,with no additional benefit of triple treatment with any of the two metabolic drugs.

**Figure S9. Whole genome sequencing based karyotypes. A.** WiDr **B**. WiDr PTPN11 KO. Estimated copy number over the different chromosomes based on GC corrected read depth. These are copy number profiles per chromosome. They show that these cell lines are similar and match with previous studies.

## Supplementary Methods

### Antibodies and reagents

WiDr cells were purchased from American Type Culture Collection (ATCC) (Prahallad *et al*, 2012) and WiDr cells clone #B32 were used as knockout of PTPN11 (WiDr PTPN11KO) (Prahallad *et al*, 2015). Both RPMI 1640 medium (#12-167F), penicillin/streptomycin (no.17-602E) and L-Glutamin (#17-605C) were purchased from Lonza, Basel, Switzerland, whereas fetal bovine serum (FBS) (#16000044) was purchased from ThermoFisher, Waltham, USA. PLX4032 (#S1267), gefitinib (#S1025) and sapitinib (#S2192) were purchased from Selleck Chemicals, Houston, TX, USA and lapatinib (#S1028) from MedKoo Bioscences, Inc. Chapel Hill, NC, USA. Etomoxir (#11969) was purchased from Bio-Connect B.V., Begonialaan 3a 6851, TE Huissen, Netherlands, whereas dichloroacetic acid sodium salt (DCA) (#2156-56-1) from Sigma-Aldrich Chemie N.V., Zwijndrecht, Netherlands.

Antibodies against HSP-90 (H-114), PTPN11 (SH-PTP2 C-18), ERK1 (C-16), ERK2 (C-14) and p-ERK1/2 (T2012/Y204, E4) were purchased from Santa Cruz. p-EGFR (Y1068, ab5644), p-SHP2 (Y542, ab62322) were purchased from Abcam. p-ERBB3 (Y1197, #4561), p-IGF1R (Y1135/1136, #3024), IGF1R (#3027), p90RSK (#8408), AKT 1/2 (#2920), p-AKT (S473, #4060)were purchased from Cell Signalling Technology. Anti-EGFR (#06-847), p-ERBB2 (Y1248, #06-229), ERBB2 (#OP15L), ERBB3 (#05-390) and p-p90RSK (T359/S363, #04-419) antibodies were from Millipore.

### Cell lines and culture conditions

Both WiDr and WiDr PTPN11 KO cells were plated in 15-cm dishes for (phospho)proteomics and 10-cm dishes for transcriptomics. All cells were cultured in RPMI supplemented with 10 % fetal calf serum (FCS) 1 % L-Glutamine and 1 % Penicillin/Streptomycin. Cells were grown around 70-80 % confluence and starved for 24 h in serum-free media, after which all plates were supplemented with serum-free media containing either no drugs (WiDr control, WiDr PTPN11 KO control) or 3 μM PLX4032 (BRAF*i,* BRAF*i* in PTPN11 KO) or 3 μM gefitinib (EGFR*i*) or the combination of 3 μM PLX4032 and 3 μM gefitinib (BRAF*i*+EGFR*i*). Following 30 min incubation, all cells were stimulated by 10 % FCS with the exception of WiDr control and WiDr PTPN11 KO control at T=0 h. Next, both untreated and treated cells were grown for 2, 6, 24 and 48 h. The whole experiment was executed in three biological replicates.

### Cell lysis

After each treatment, both WiDr and WiDr PTPN11 KO cells were harvested by washing twice with cold PBS and then resuspended in ice cold lysis buffer. For (phospho)proteomics analysis, protein extraction was obtained adding 2 mL of buffer containing 8 M urea, 50 mM ammonium bicarbonate (pH 8.0), 1 mM sodium orthovanadate, complete EDTA-free protease inhibitor mixture (Roche), and phosSTOP phosphatase inhibitor mixture (Roche) was used. Finally cells were snap frozen in 15 mL Falcon Centrifuge Tubes and stored at -80 °C until use. For transcriptomics analysis, mRNA extraction was achieved adding 600 μL Lysis/Binding Buffer from the *mir*Vana miRNA Isolation Kit by Ambion (Cat. AM1560), and then collecting cells by scraping. Next, cell lysate were transferred into a 1.5 mL tube and stored at -80 °C until time of total RNA isolation.

### Transcriptomics sequencing

Quality and quantity of isolated RNA was checked and measured with Agilent 2100 Bioanalyzer and RNA Nano 6000 chips (Agilent, Cat. 5067-1511). Library were generated from 500 ng of Total RNA using the Truseq Stranded Total RNA kit with Ribo-Zero Human/Mouse/Rat set A and B by Illumina (Cat. RS-122-2201 and RS-122-2202). After the library preparation, libraries were checked with Bioanalyzer2100 DNA High Sensitivity chips (Cat. 5067-4626) and with Qubit (Qubit^®^ dsDNA HS Assay Kit, Cat. Q32854). Libraries were equimolar pooled to 2 nM. Next, 1.0-1.4 pM of these pooled libraries were sequenced on the Illumina NextSeq, 2x75bp high output, and 1.0-1.4 pM of library pools was loaded. Mapping was performed using STAR_2.4.2a, read counting using ht-seq count and the v74 gencode definition for coding regions. Fragments were mapped against GRCh37 using STAR (Dobin *et al*, 2013) (v 2.4.2a), reads within coding regions (ENSEMBL release 74) were counted using ht-seq count (Anders *et al*, 2015) (v0.6.1) and further normalized and analysed using the DeSeq2 package (Kim *et al*, 2013) (v1.6.3). Full details and workflows are available online: https://github.com/UMCUGenetics/RNASeq (v.2.2.0 was used for this paper).

### Proteomics analyses

Lysed cells were defrost and each tube was supplemented with 300 μL of fresh lysis buffer. Cells were further lysed by 10 rapid passages through 23G needle and by sonication on ice. Cell debris were removed by centrifugation at 20,000 x *g* for 30 min at 4 °C and cleared supernatants were stored at -80 °C. The total protein concentration was measured using Bradford assay (Bio-Rad).

Next, samples were split into 200 μg aliquots for quality control analysis via western blots and 2 mg aliquots for tryptic digestion. Proteins were reduced with 8 mM DTT for 1 h at room temperature, alkylated with 16 mM iodoacetamide for 30 min at room temperature in the dark and reduced again with 8 mM DTT at room temperature to prevent overalkylation. Later, proteins were first digested by Lys-C (enzyme/substrate ratio 1:65) at 37 °C for 4 h. Subsequently, urea was diluted to 2 M with 50 mM ammonium bicarbonate and trypsin was added (enzyme/substrate ratio 1:50). The digestion was executed at 37 °C overnight and then quenched with 5 % formic acid. Peptides were desalted using Sep-Pak C18 cartridges (Waters), dried and stored at -80 °C.

### Western Blots

For both quality control and validation analysis, total cell extracts were quantified using the Pierce BCA Protein Assay (23227, Thermo Scientific) and the colorimetric reaction evaluated at 562 nm using the EnVision 2014 Microplate Reader (Perkin Elmer). Equal amount of proteins were prepared for all samples adding 10X NuPage Sample Reducing Agent (NP0004, Thermo Scientific) and 4X NuPage LDS Sample Buffer (NP0007, Thermo Scientific). Samples were subsequently incubated at 95 °C for 5 min to allow protein denaturation. Lysates were resolved by SDS-PAGE using NuPage 4-12% Bis-Tris precast gels and NuPAGE Gel Electrophoresis Systems (Thermo Scientific). The gels were run in 1X MOPS buffer (50 mM MOPS, 50 mM Tris base, 0.1 % SDS, 1 mM EDTA) at a constant voltage of 165 V. Proteins were transferred on a methanol-activated PVDF membrane. Transfer was performed in 1X Transfer Buffer (25 mM Tris base, 122 mM glycine, 0.01 % SDS, 10 % methanol) using a Trans-Blot Cell apparatus (Bio-Rad) and applying a constant amperage of 70 mA. Blocking was performed by incubating the membranes in 5 % BSA in TBS-T (0.1 %) for 1 h. Primary antibodies were typically diluted 1:1000 in 5% BSA in TBS-T and incubated at 4 °C overnight while shaking. Membranes were washed 3 times during 10 min with TBS-T (0.1 %). HRP-coniugated secondary antibodies (Bio-Rad) were diluted 1:10,000 in 5% BSA in TBS-T and incubated for 1 h at room temperature while shaking. Subsequently, membranes were washed additional 3 times during 10 min with TBS-T (0.1 %). Final protein detection was performed using Clarity ECL Western Blotting substrate (Bio-Rad) and blot imaging was performed using the Chemidoc Touch Imaging System (Bio-Rad).

### Phosphopeptide enrichment

Ti^4+^-IMAC material was prepared and used as previously described(Zhou *et al*, 2013). Briefly, the affinity material was loaded onto GELoader tips (Eppendorf) using a C_8_ plug. The columns were pre-equilibrated two times with 50 μL of Ti^4+^-IMAC loading buffer (80 % ACN, 6 % trifluoroacetic acid (TFA)). Next, samples were resuspended in loading buffer and 200 μg were loaded into each microcolumn. Columns were sequentially washed with 50 μL wash buffer A (50% ACN, 0.5 % TFA, 200 mM NaCl) and 50 μL wash buffer B (50 % ACN, 0.1 % TFA). Bound peptides were first eluted by 30 μL of 10 % ammonia into 30 μLof 10 % FA. Finally, all remaining peptides were eluted with 2 μL of 80 % ACN, 2 % FA. The collected eluate was further acidified by adding 3 μL of 100 % FA, and subsequently dried in *vacuo* and stored at -80°C. The procedure was repeated in three technical replicates for each biological replicate. Later, phosphopeptides were further desalted using SPE C18 cartridge homemade. Stationary phase C_18_ beads were dissolved in 500 μL isopropanol and loaded onto GELoader tips (Eppendorf) using a C_18_ plug as previously described. The columns were washed with 50 μL wash buffer C (80 % ACN, 0.1 % TFA) and then conditioned with 0.1 % TFA. Next, samples were resuspended in 30 μL of 10 % TFA and loaded into columns which were further washed with 30 μL of 10 % TFA. Finally, phosphopeptides were eluted with 30 μL of 80 % ACN, 1 % FA, dried and stored at -80 °C.

### (Phospho)proteomics mass spectrometry

(Phospho)peptides were analysed using an Agilent 1290 Infinity II LC system coupled to a Q-Exactive mass spectrometer (Thermo Scientific). The LC system was equipped with a 2 cm Aqua C18 (Phenomenex) trapping column (packed in-house, i.d., 50 μm; resin 5 μm) and a 40 cm Poroshell 120 EC-C18 (Agilent) analytical column (packed in-house, i.d. 50 μm; resin 3 μm). (Phospho)peptides were first trapped at 5 μL/min in 100 % solvent A (0.1 % formic acid in water) for 10 min, and then eluted with solvent B (100 % ACN/0.1 % formic acid) at a flow rate of 200 μL/min. Phoshoproteome analysis was performed in 120 min gradient as follows: 0-10 min 100 % solvent A, 10–105 min 4 % solvent B, 105-108 min 36 % solvent B, 108-109 min 100% B, 109-120 min 100 % solvent A. For the proteome analysis instead a 180 min gradient was set as follows: 0-10 min 100 % solvent A, 10–10.1 min 13 % solvent B, 10.1-165 min 40 % solvent B, 165-168 min 100 % B, 169-180 min 100 % solvent A. The electrospray voltage was set to 1.7 kV using a coated SilicaTip P200P capillary (Thermo Scientific). The mass spectrometer was operated in data-dependent acquisition mode and was configured to perform a Fourier transform survey scan from 375 to 1600 *m/z* (resolution 35,000) followed by higher collision energy dissociation (HCD) fragmentation of the 10 most intense peaks (25 % normalized collision energy at a target value of 50,000 ions, resolution 17,500). In total, 411 RAW spectra were collected: 168 for proteomics and 243 for phosphoproteomics.

### (Phospho)proteomics data processing

A preliminary quality analysis was performed for each dataset by MaxQuant (version 1.5.1.2) using a label-free quantification approach and Swissprot database *Homo sapiens* database (write the number of entries and possibly the version). A cut-off of R^2^=0.7 was applied on both proteomics and phosphoproteomics correlation matrix (data not shown) and, after this filtering step, 383 RAW phosphoproteomics and proteomics files were selected for the integrated analysis and analyzed by MaxQuant (version 1.5.2.8). Phosphoproteomics RAW data were classified as Group 0, whereas proteomics RAW data as Group 1. A mutant-modified Swissprot database *Homo sapiens* (24,126 entries, released on 05_2015) was used for the database search: for the variant identification, all STAR aligned reads were merged into a single bam file, subsequent variant calling was performed using VarScan (Koboldt *et al*, 2009) (v2.3.8) with default settings; resulting variant positions were included in the extended database when covered by at least 100 reads with at least 20 % of reads harboring the variant. Trypsin was specified as enzyme and up to two missed cleavages were allowed. Cysteine carbamidomethylation was set as a fixed modification, while methionine oxidation and protein N-term acetylation were set as variable modifications. Phosphorylation on serine, threonine and tyrosine was also selected as variable modification for the phosphoproteomics analysis. The allowed fragment mass deviation was set to 20 ppm for FTMS, and both minimum peptide length and the maximum peptide charge were set to 7. Fast Label free quantification (LFQ) was performed and ‘match between runs’ was enabled. Peptide and protein identification was set to 1 % FDR. The quantified output (*proteinGroups.tx*t, *phospho (STY)Sites.txt*) were processed using a custom Python package (PaDuA). Potential contaminants and reverse peptides were removed, and filters for Class I phosphosites (localization probability > 75 %) and ‘only identified by site’ were applied for phospho-and protein data respectively. Normalization was performed by subtracting the median of log_2_ transformed intensities from each column. For phospho-data, *‘*expand side table’ function was applied before normalization. Median of technical replicates was performed for each dataset and resulting values were filtered to ensure each protein or phosphosite had valid measurements in at least one time point of any of the six cell culture conditions. Enrichment analysis was calculated using *modificationSpecificPeptides.txt* table. Final processed output were exported for subsequent analysis in R.

### Clustering

Hierarchical clustering was performed on the 1500 phosphosites, proteins, or mRNAs with the highest variance within each dataset, calculated over all conditions. The mRNA expression data was first log-transformed and filtered for mRNAs that were differentially expressed compared to the T=0 h control at a false discovery rate (FDR) of 10^-6^. The pairwise distance between two phosphosites, proteins, or mRNAs i and j was calculated as (1-ρ_i,j_)/2, where ρ_i,j_ is the Pearson correlation between the two. The obtained distances were used for hierarchical clustering using Ward’s minimum variance method. For each data type, the resulting trees were cut into 8 groups, since consensus clustering (Wilkerson & Hayes, 2010) indicated that to be a reasonable number. Enrichment analysis of the clusters were done using Fisher’s exact test, with all measured genes, proteins, or phosphosites as background. For the phosphoproteomic data, the enrichment for each kinase in each cluster was calculated based on its predicted substrates. Predicted kinase-substrate relations were obtained using KinomeXplorer (Horn *et al*, 2014). Similarly, for the mRNA expression data, enrichment of transcription factors in each cluster was calculated based on its known target genes. Transcription factor-target gene relations were obtained from the TransFac database (Matys *et al*, 2006; Wingender *et al*, 1996) (version 2_2016). Enrichment analysis of biological processes was done using the hallmarks gene sets from MSigDB (Liberzon *et al*, 2015). Multiple testing correction was done using Benjamini and Hochberg’s method.

### Cell proliferation assays

All experiments were carried out culturing WiDr cells in RPMI 1640 supplemented with 10 % fetal bovine serum (FBS), 1 % Penicillin/streptomycin and 1 % L-Glutamine, at 37 °C, 5 % CO_2_. WiDr cells were seeded in five 96-well plates at a density of 5000 cells/well. After 24 hours of incubation (37 °C, 5 % CO_2_), media was removed from all plates and replaced with media containing drugs. In three 96 wells plates, gefitinib, lapatinib or sapitinib were serially diluted to a final concentration range of 120 nM to 30 μM as a single treatment and in combination with PLX4032 at a fixed concentration of 3 μM (n=4). In the fourth 96-well plate, PLX4032 was serially diluted to a final concentration range of 30 nM to 30 μM in combination with gefitinib at a fixed concentration of 3 μM and with either 1 mM DCA (n=4) or 10 μM etomoxir (n=4) respectively. In the last 96-well plate, PLX4032 was serially diluted to a final concentration range of 30 nM to 30 μM in combination with gefitinib at a fixed concentration of 3 μM (n=8). After 96 h of treatment, 1 mM DCA was added to four replicates and 10 μM etomoxir was added to other four replicates. All plates contained a column with untreated cells as a reference sample and were treated until 7 days (37 °C, 5 % CO_2_). Media containing the drugs was replaced after 72 h.

IC50 of single treatments was determined seeding WiDr cells in three 96-well plates at a density of 10,000 cells/well. After 24 hours of incubation (37 °C, 5 % CO_2_), media was removed from all plates and replaced with media containing either PLX4032 or etomoxir or DCA serially diluted in four replicates to a final concentration range 30 nM-30 μM, 200 μM-0.2 μM and 100 mM-0.1 mM, respectively. All plates contained a column with untreated cells as a reference sample and were treated until 72 h (37 °C, 5 % CO_2_).

Cell growth inhibition was monitored in all assays using the IncuCyte™ automated microscope (Essen Bioscience, Ann Arbor, USA) and phase-contrast images were collected every 2 h using a 10x Nikon objective. Phase confluence percentage from each well at each time point was exported into GraphPad Prism 7.0 software. The area under the curve (AUC) was calculated for each concentration (n=4), normalized in respect to untreated cells and fitted using a four-parameter logistic curve. Percentage of growth in single and combination treatments were visualized as dose response curves.

### Drug off assays

All experiments were carried out culturing WiDr cells in RPMI 1640, supplemented with 10 % fetal bovine serum (FBS), 1 % Penicillin/streptomycin and 1 % L-Glutamine at 37 °C, 5 % CO_2_. For long-term assay, WiDr cells were seeded in one 6-well plate (200,000 cells/well) and grown to around 60 % confluence. After 24 h starvation in serum-free media, cells were treated with PLX4032 and gefitinib both at a fixed concentration of 3 μM, in complete medium. Media was replaced two times per week and treatment was interrupted after 78 days. Pictures were acquired using an ECLIPSE Ti-e inverted microscope (Nikon, Tokyo, Japan) at a magnification of 10x. Cell confluence was measured using Fiji plugin in ImageJ software.

For short-term assay, WiDr cells were seeded in one 96-well plate at a density of 5000 cells/well. After 24 hours of incubation (37 °C, 5 % CO2), media was removed from all plates and replaced with media containing PLX4032 and gefitinib at fixed concentration of 3 μM. Media was replaced (once/twice) per week and treatment was interrupted after 5 days. Cell growth was monitored in the IncuCyte™ automated microscope (Essen Bioscience, Ann Arbor, USA) and phase-contrast images were collected every 2 h using objective Nikon 10x.

